# Selective activation of subthalamic nucleus output quantitatively scales movements

**DOI:** 10.1101/2022.01.19.477002

**Authors:** Alexander D. Friedman, Henry H. Yin

## Abstract

The subthalamic nucleus (STN), a diencephalic nucleus strongly connected with the basal ganglia, is a common target for deep brain stimulation (DBS) treatments of Parkinsonian motor symptoms. The STN is thought to suppress movement by enhancing inhibitory basal ganglia output via the indirect pathway, and disrupting STN output using DBS can restore movement in Parkinson’s patients. However, previous studies on STN DBS usually relied on electrical stimulation, which cannot selectively target STN output neurons. Moreover, most studies did not measure behavior precisely during STN manipulations. Here we selectively stimulated STN projection neurons, by expressing channelrhodopsin in vesicular glutamate transporter 2 (VGlut2)-positive projection neurons in the STN. We also quantified behavior in freely moving male mice with high spatial and temporal resolution using 3D motion capture at 200 frames per second. We found STN stimulation resulted in movements with very short latencies (10-15 ms). Unilateral stimulation quantitatively determines orientation, head yaw, and head roll consistently in the ipsiversive direction, toward thee side of stimulation. On the other hand, head raising (increased head pitch) was observed after stimulation on either side. Strikingly, a single pulse of light was sufficient to generate movement, and there was a highly linear relationship between stimulation frequency and kinematics. These results question the common assumption that STN suppresses movement; instead they suggest that STN output can precisely specific action parameters via direct projections to brainstem regions.

## Introduction

The subthalamic nucleus (STN) is a key diencephalic nucleus that is strongly and reciprocally connected with the basal ganglia (BG)(Parent and Hazrati, 1995). It is sometimes considered to be a part of the indirect pathway in the BG. According to classic models, the direct and indirect pathways play opposing roles, with the former responsible for movement initiation and the latter for movement inhibition (Albin et al., 1989; Kravitz et al., 2010). The direct pathway exerts its influence through inhibition of BG output nuclei such as internal globus pallidus (GPi) and substantia nigra pars reticulata (SNr), whereas the indirect pathway can disinhibit the GPi and SNr, in part via the STN (Smith and Bolam, 1989; Alexander and Crutcher, 1990; DeLong, 1990). Given the STN’s glutamatergic projections to the GPi, its activation is thought to oppose the effects of the direct pathway, resulting in movement cancelation (Nambu et al., 2002; Aron and Poldrack, 2006; Fife et al., 2017). Consequently, STN is often used as the site of deep brain stimulation (DBS), a major treatment option for Parkinson’s Disease (PD), as STN stimulation at high frequency is thought to reduce hyperactive indirect pathway activity caused by DA depletion (DeLong and Wichmann, 2001).

However, evidence for the STN’s role in canceling action is mixed. Although human fMRI work using a stop-signal task suggested that the STN is more activated during successful stop compared to go trials (Aron and Poldrack, 2006), electrophysiological results in rats using a similar task showed that STN neurons increase firing after stop cues regardless of whether stopping was successful (Schmidt et al., 2013). There have also been conflicting observations on the effects of STN stimulation on movement. Bilateral STN photo-stimulation was reported to reduce horizontal movement while inhibition increased horizontal movement, but the opposite pattern was observed in grooming behavior (Guillaumin et al., 2021). Moreover, in another study, STN stimulation failed to cause a significant effect on locomotion in the open field altogether (Heston et al., 2020). Bilateral lesions of the rat STN have also been shown to affect stopping accuracy in a stop-signal task, but stop signal reaction time was unaffected, and the stopping deficit did not depend on the stop-signal delay, suggesting that lesions may actually have increased impulsive behavior rather than simply caused problems stopping behavior (Eagle et al., 2008). Indeed, impulsive responding has been found with bilateral rat STN lesions in several tasks (Baunez et al., 1995; Baunez and Robbins, 1997; Uslaner and Robinson, 2006).

Fife et al. (2017) reported that photo-stimulation of the mouse STN cancels ongoing lick bouts or induces a pause when bouts are not canceled, but video evidence suggests stimulation actually caused head and body movements that interfered with licking. Although there was some data indicating that STN activation increases head movement and turning behavior, the active role of the STN in movement has not been examined closely (Heston et al., 2020; Guillaumin et al., 2021).

A major limitation of previous work is the lack of precise quantification of the movements during STN manipulations. In the present study, we determined the relationship between photo-stimulation parameters and movement measured with high temporal and spatial resolution. In particular, 3D motion capture measuring movement at 200 frames per second was used to quantify the subtle changes in head and body angle caused by optogenetic stimulation. To selectively target STN projection neurons, we used a vGlut2-Cre driver line combined with a Cre-dependent viral vector with channelrhodopsin. We found that selective STN activation generates movement of the head and torso with very short latency, and that there was a highly linear relationship between stimulation frequency and movement kinematics.

## Materials and Methods

### Subjects

Data was collected from 11 mice. The experimental group consisted of 7 adult male Vglut2-ires-Cre mice (Slc17a6^tm2(cre)Lowl^, Jackson Labs, ME), which express Cre-recombinase under control of Vglut2 receptor regulatory elements. These mice enabled selective expression of Cre-dependent channelrhodopsin (ChR2) in the STN, because surrounding areas are devoid of VGlut2. The control group consisted of 4 adult (2 males, 2 females) wild type mice. Mice were housed in groups of 2-5 and given free access to food and water. They were maintained on a 12:12 light/dark cycle, and experiments were conducted during the light phase. All experimental procedures were approved by the Duke University Institutional Animal Care and Use Committee.

### Surgery

Mice were anesthetized with 3% isoflurane in oxygen flowing at a rate of 0.6 L/min. They were then secured in a stereotactic frame (David Kopf Instruments, CA) and injected with Meloxicam (2 mg/kg) before incision. Next, 150 to 400 nl of a viral vector containing a Cre-dependent ChR2 (pAAV5.EF1a.DIO.hChR2(E123T/T159C).eYFP, Addgene) was injected bilaterally into the STN (AP: −2.0, ML: ±1.6, DV: −4.5 mm relative to bregma) at 1 nl/s with a Nanoject III microinjector (Drummond Scientific, PA). Custom-made optic fibers (0.22NA, MM, 105 μm core, >80% transmittance, 5 mm below ferrule) were then lowered directly into the injection tract and secured 150 μm above the injection site with dental acrylic anchored to skull screws (Figure 1A). Custom-made headbars (H.E. Parmer, TN) were then affixed to the acrylic. Mice were given 4 weeks to recover from surgery before testing.

**Figure 1.**
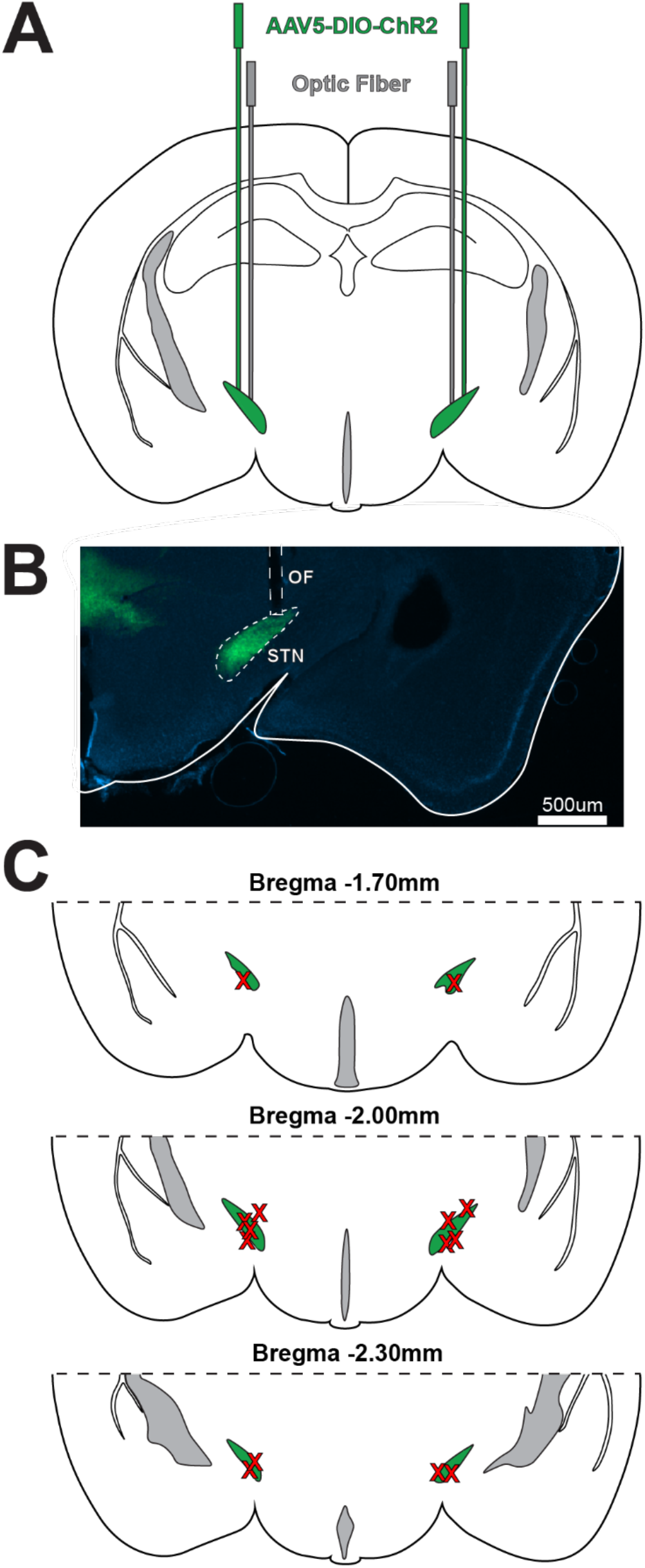
Placement of viral injections and optic fibers. (A) Schematic of viral injection and fiber optic implantation strategy in bilateral STN of VGlut2-Cre mice. (B) Representative image of eYFP expression in STN of VGlut2-Cre mouse injected with AAV5-DIO-ChR2-eYFP. (C) Optic fiber tip locations.

### Open Field Stimulation & 3D Motion Capture

Mice were placed on a one foot tall, wall-less, 8×8 inch open field platform, set on a table centrally located in the experiment room. 6.4mm diameter infrared markers (B&L Engineering, CA) were attached just above and to either side of the head, at the base of the optic fibers and on the headbars, with another added to the tail base. 6 Osprey motion tracking cameras (Motion Analysis, CA) positioned around the table captured marker positions at 200 Hz, and the Cortex Motion Analysis computer program saved them in 3D cartesian coordinates (Figure 2A) (Bartholomew et al., 2016).

**Figure 2.**
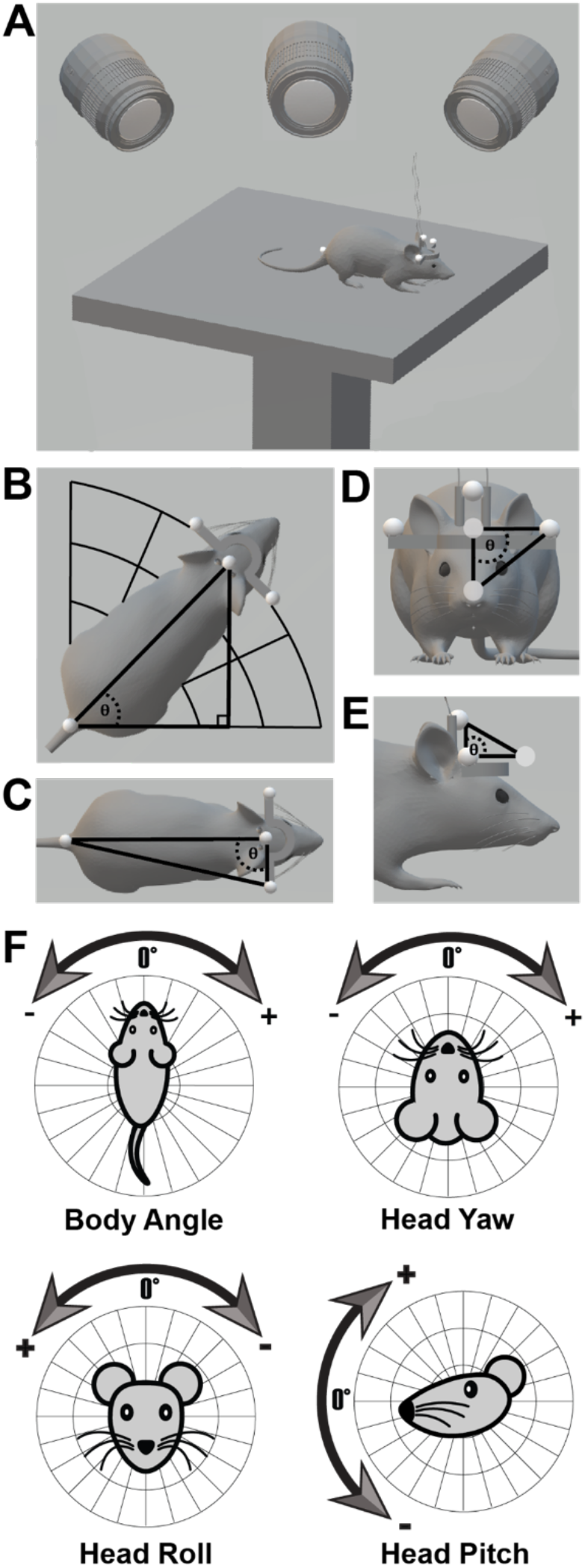
Quantifying movement kinematics using 3D motion capture. (A) Schematic of the experimental setup. (B) Body angle calculation. A right triangle was made in the X-Y plane with the tail marker and center of the left and right head markers (head center) as vertices. The angle at the tail marker was shifted into the proper polar quadrant based on the mouse’s orientation and was used as body angle. (C) to calculate yaw triangle was made in the X-Y plane with the tail marker, right head marker, and head center as vertices. The angle at head center minus 90 was used as yaw. (D) To compute roll, a triangle was made in the X-Z plane with the left head marker, head center, and a float point 10 mm below head center as vertices. The angle at head center minus 90 was used as roll. (E) To calculate pitch, a triangle was made in 3D space with the top head marker, head center, and a float point 10 mm in front of head center as vertices. The angle at head center minus 90 was used as pitch. (F) Illustration of the key measures: body angle, head yaw, roll, and pitch.

MATLAB (MathWorks, MA) was used to make a NI USB-6211 (National Instruments, TX) operate a diode-pumped solid state laser (Shanghai Laser & Optics Century Co., Ltd., China) at specific frequencies and pulse widths, detailed below. Laser light was relayed through a 100 μm core optic fiber (Precision Fiber Products, CA - 0.22NA, MM), to a FRJ_1×2i_FC-2FC_0.22 commutator (Doric Lenses, Canada), into another 100 μm core optic fiber (Precision Fiber Products, CA - 0.22 NA, MM), which was adjusted to output 7-12 mW on a mouse by mouse basis. The fiber was then connected to the mice’s fiber optic implants. A Cerebus Neural Signal Processor (Blackrock Microsystems, UT) was used to record the laser control signal from the NI device and the frame signal from the Osprey cameras, so movement could be aligned to laser pulses.

All single pulse data from a given STN was collected in a single session. 5, 10, 20, and 40 ms pulses were delivered pseudo-randomly, so that each occurred 15 times, with a 15-20s inter-pulse interval. All frequency data from a given STN was collected in a separate single session. Half second, 10 ms pulse width, 5, 10, 20, and 40 Hz laser trains were delivered pseudo-randomly, so that each occurred 15 times, with a 20-30s inter-train interval. Data from a given mouse was collected in a single day, with single pulse sessions preceding frequency sessions.

### Analysis and Statistics

Calculation of body angle, head yaw, head roll, and head pitch was done in MATLAB for each motion sample. The MATLAB function “unwrap” was used to remove discontinuities caused by the body angle changing from 0 to 360 degrees, or vice versa (Figure 2B). Velocity was calculated for each variable by finding the difference in position across adjoining samples and dividing by the inverse frame rate.

Movement following a stimulation train was calculated for each combination of mouse, behavioral variable, STN, and frequency. Movement following single laser pulses was quantified for each combination of mouse, behavioral variable, STN, and pulse width. Latency to move following stimulation onset was determined for each combination of mouse, behavioral variable, and STN by manual inspection of velocity changes caused by 40 ms single pulse stimulation. The time of the first significant change from pre-stimulation baseline in the effect direction was chosen on trials in which such a change occurred before stimulation offset, and the results were averaged across trials. Velocity 60 and 120 ms after single laser pulse onset was calculated for each combination of mouse, behavioral variable, STN, and pulse width by finding the velocity 60 and 120 ms after each laser pulse onset and averaging across each timepoint.

All statistical tests (linear regression, ANOVAs, t-tests) were performed in GraphPad Prism (GraphPad Software, San Diego, CA).

### Histology

Mice were perfused with 0.1M phosphate buffered saline followed by 4% paraformaldehyde. Brains were then stored in 4% paraformaldehyde with 30% sucrose for 72 hours, before being transferred to 30% sucrose for another 24. Brains were then sliced into 60 μm coronal sections with a Leica CM1850 cryostat. After, sections were mounted and immediately coverslipped with Fluoromount-G with DAPI medium (catalog no. 17984-24, Electron Microscopy Sciences). To confirm viral expression and fiber placement, slices were examined with an Axio Zoom V16 microscope (Zeiss, NY) (Figure 1B-C).

## Results

VGlut2-Cre and WT control mice injected with Cre-dependent ChR2 were stimulated for 500 ms, unilaterally in each STN, at 5, 10, 20, and 40 Hz. First, body angle changes as a result of stimulation were quantified. Unilateral stimulation revealed a linear relationship between stimulation frequency and turning angle (linear regression: r^2^ = 0.9993, p = 0.0004; r^2^ = 0.9922, p = 0.0039; Figure 3A, Left). There was also a significant interaction between side of stimulation and genotype on body turning caused by 40 Hz stimulation, which generated the largest effect (2-Way ANOVA: F(1,18) = 23.36, p = 0.0001). Left STN stimulation caused significant leftwards body turning compared with controls (p = 0.0035) and right STN stimulation (p < 0.0001). Right STN stimulation caused significant rightwards body turning compared to controls (p = 0.0108) or left STN stimulation (p < 0.0001) (Figure 3A, Right).

**Figure 3.**
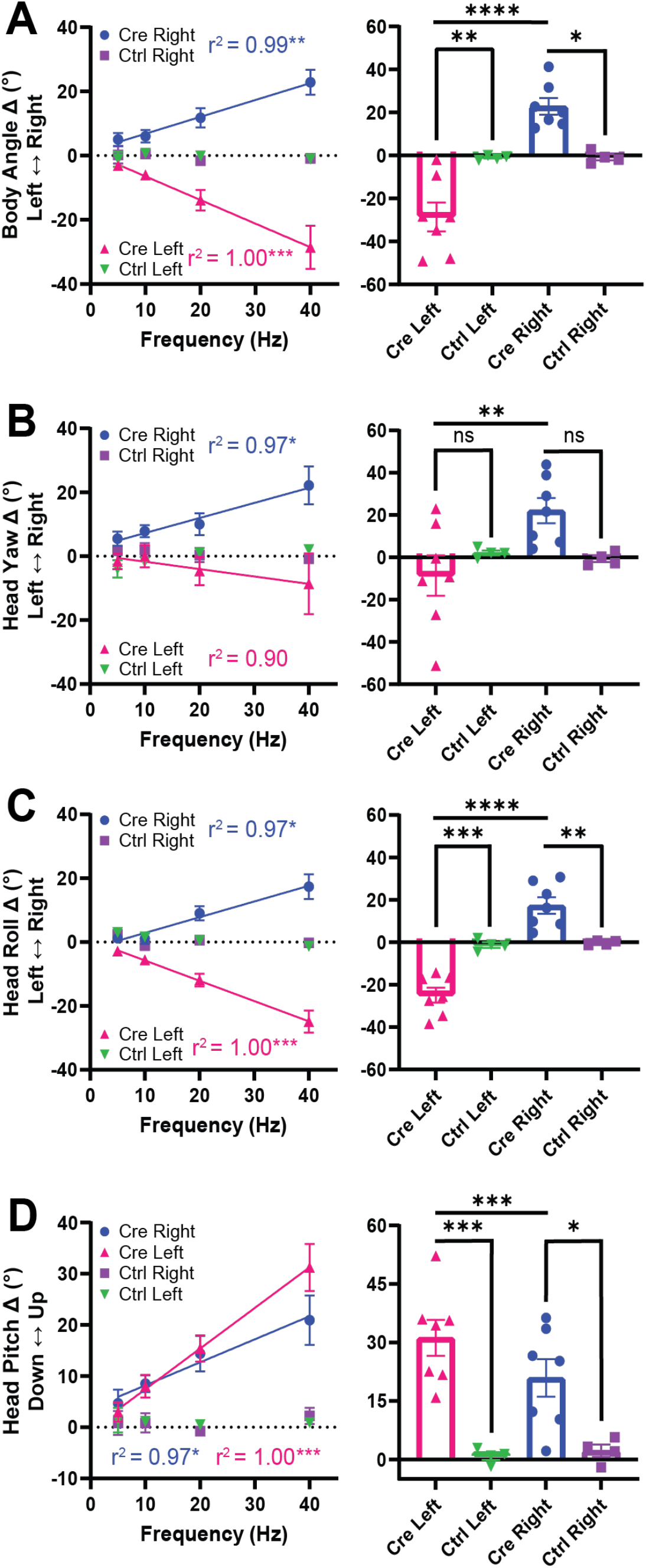
Photo-stimulation of STN VGlut2+ neurons causes movement (frequency manipulation). (A) *Left:* Linear relationships between stimulation frequency and ipsiversive turning with left and right STN stimulation. *Right:* Significant interaction between STN and genotype on turning caused by 40 Hz stimulation (2-Way ANOVA). Left and right STN stimulation in Cre mice causes significant ipsiversive turning vs. controls. (B) *Left:* Linear relationships between stimulation frequency and ipsiversive head yaw with left and right STN stimulation. *Right:* Significant interaction between STN and genotype on head yaw caused by 40Hz stimulation (2-Way ANOVA). Left and right STN stimulation causes significant ipsiversive head yaw when compared within Cre mice. (C) *Left:* Linear relationships between stimulation frequency and ipsiversive head roll with left and right STN stimulation. *Right:* Significant interaction between STN and genotype on head roll caused by 40Hz stimulation (2-Way ANOVA). Left and right STN stimulation in Cre mice causes significant ipsiversive head roll vs. controls. (D) *Left:* Linear relationships between frequency and upwards head pitch with left and right STN stimulation. *Right:* Significant interaction between STN and genotype on head pitch caused by 40Hz stimulation (2-Way ANOVA). Left and right STN stimulation in Cre mice causes significant upwards head pitch vs. controls, with a stronger effect of stimulation in the left vs. right STN. *p<0.05, **p<0.01, ***p<0.001, ****p<0.0001. Error bars indicate SEM.

In VGlut2-Cre mice, unilateral stimulation revealed a linear relationship between frequency and yaw (linear regression leftward: r^2^ = 0.8976, p = 0.0526; rightward: r^2^ = 0.9693, p = 0.0155) (Figure 3B, Left). There was also a significant interaction between stimulation side and genotype on head yaw caused by 40Hz stimulation (2-Way ANOVA: F(1,18) = 4.824, p = 0.0414). Left STN stimulation caused significant leftwards head yaw versus right STN stimulation (p = 0.0074). Right STN stimulation caused significant rightwards head yaw versus left STN stimulation (p = 0.0074) and a trending effect versus right STN stimulation in controls (p = 0.0949). Head roll changes due to stimulation were also measured. Unilateral stimulation revealed a linear relationships between stimulation frequency and head roll (linear regression: F(1,2) = 3534, p = 0.0003; r^2^ = 0.9994; linear regression: F(1,2) = 63.57, p = 0.0154; r^2^ = 0.9695) (Figure 3C, Left). There was also a significant interaction between stimulation side and genotype on head roll caused by 40Hz stimulation (2-Way ANOVA: F(1,18) = 32.76, p < 0.0001). Left STN stimulation caused significant ipsiversive roll compared to controls (p = 0.0004) and right STN stimulation (p < 0.0001). Right STN stimulation caused significant ipsiversive roll compared to controls (p = 0.0057) and left STN stimulation (p < 0.0001) (Figure 3C, Right).

Unilateral stimulation revealed a significant linear relationship between frequency and upwards head pitch (left, linear regression: r^2^ = 0.9987, p = 0.0006; right, linear regression: r^2^ = 0.9652, p = 0.0176) (Figure 3D, Left). There was also a significant interaction between STN and genotype on head pitch caused by 40 Hz stimulation (2-Way ANOVA: F(1,9) = 21.77, p = 0.0012). Left and right STN stimulation caused upwards head pitch versus stimulation in the left and right STNs of controls (Sidak’s test, p = 0.0004, p = 0.0197), but the effect was greater in the left versus right STN (p = 0.0002) (Figure 3D, Right).

The same mice were also stimulated unilaterally in each STN with single laser pulses while pulse width was manipulated. First, mean body angle changes 100 ms after stimulation onset were measured. Left and right STN stimulation revealed significant linear relationships between laser pulse width and body turning (linear regression: F(1,2) = 101.5, p = 0.0097; r^2^ = 0.9807; linear regression: F(1,2) = 35.33, p = 0.0272; r^2^ = 0.9464) (Figure 4A, Left). There was also a significant interaction between STN and genotype on body turning caused by 40 ms stimulation (2-Way ANOVA: F(1,18) = 19.12, p = 0.0004). Left STN stimulation caused significant leftwards body turning versus left STN stimulation in controls (p = 0.0085) and right STN stimulation (p < 0.0001). Right STN stimulation caused significant rightwards body turning versus right STN stimulation in controls (p = 0.0185) and left STN stimulation (p < 0.0001) (Figure 4A, Right). Second, mean angular changes in head yaw 100 ms after stimulation onset were measured. Right STN stimulation revealed a significant linear relationship between laser pulse width and rightwards head yaw (linear regression: F(1,2) = 72.48, p = 0.0135; r^2^ = 0.9731). However, left STN stimulation did not reveal a significant relationship between laser pulse width and head yaw (linear regression: F(1,2) = 0.4545, p = 0.5697; r^2^ = 0.1852) (Figure 4B, Left). There was also a significant interaction between STN and genotype on head yaw caused by 40 ms stimulation (2-Way ANOVA: F(1,18) = 12.12, p = 0.0027). Right STN stimulation caused significant rightwards head yaw versus right STN stimulation in controls (p = 0.0118) and left STN stimulation (p < 0.0001). Left STN stimulation caused significant leftwards head yaw versus right STN stimulation (p < 0.0001) (Figure 4B, Right). Third, mean angular changes in head roll 100 ms after stimulation onset were measured. Left and right STN stimulation revealed significant linear relationships between laser pulse width and roll of the head (linear regression: F(1,2) = 139.5, p = 0.0071; r^2^ = 0.9859; linear regression: F(1,2) = 29.00, p = 0.0328; r^2^ = 0.9355) (Figure 4C, Left). There was also a significant interaction between STN and genotype on head roll caused by 40 ms stimulation (2-Way ANOVA: F(1,18) = 13.87, p = 0.0016). Left STN stimulation caused significant leftwards head roll versus left STN stimulation in controls (p = 0.0042) and right STN stimulation (p < 0.0001). Right STN stimulation caused significant rightwards head roll (p < 0.0001, Figure 4C, Right). Changes in head pitch 100 ms after stimulation onset were also measured. Left and right STN stimulation revealed significant a linear relationship between laser pulse width and upwards head movement (linear regression: F(1,2) = 666.2, p = 0.0015; r^2^ = 0.9970; linear regression: F(1,2) = 35.39, p = 0.0271; r^2^ = 0.9465; Figure 4D, Left). There was also a significant main effect of genotype on head pitch caused by 40 ms stimulation (2-Way ANOVA: F(1,9) = 20.15, p = 0.0015; Figure 4D, Right).

**Figure 4.**
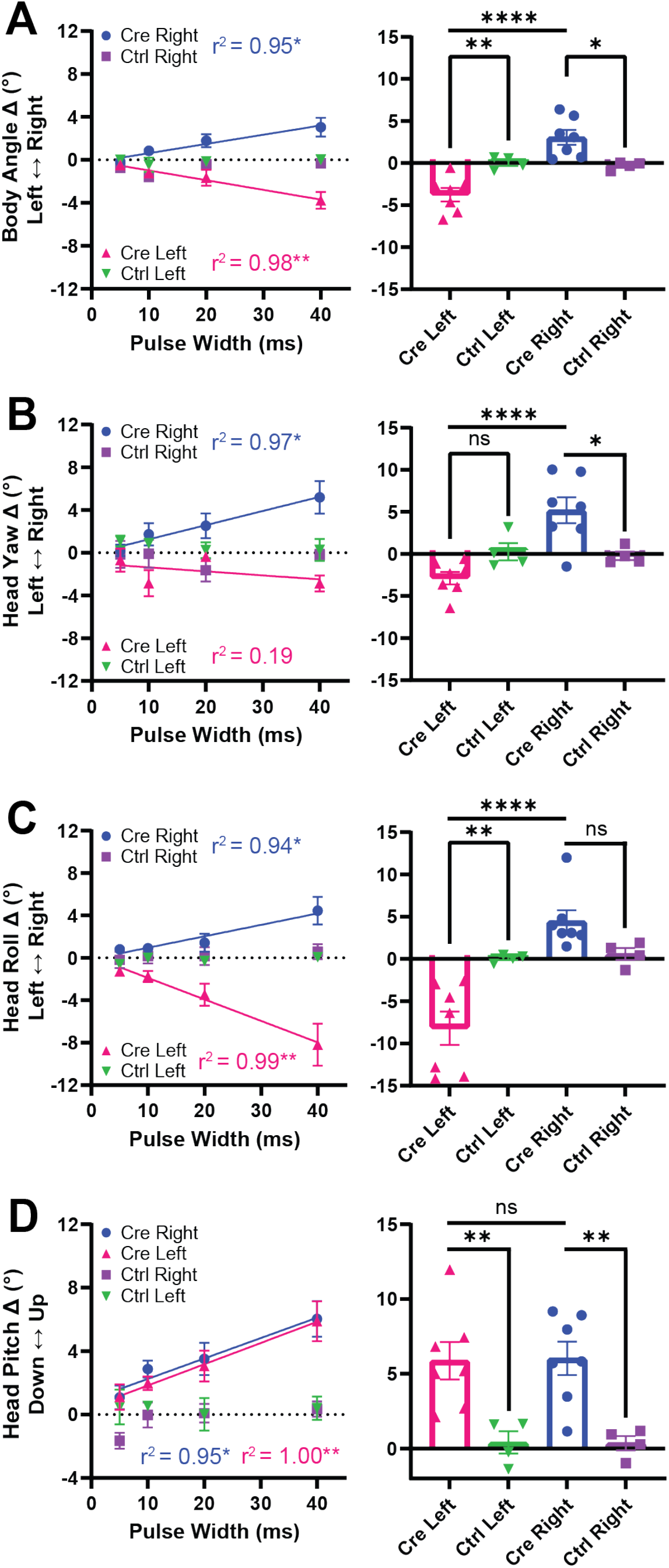
Body and head angle changes after stimulation (pulse width manipulation). (A) *Left:* Linear relationships between pulse width and ipsiversive turning with left and right STN stimulation. *Right:* Significant interaction between STN and genotype on turning caused by 40 ms stimulation (2-Way ANOVA). Left and right STN stimulation in Cre mice causes significant ipsiversive turning vs. controls. (B) *Left:* Linear relationship between pulse width and ipsiversive head yaw with right STN stimulation. No significant relationship with left STN stimulation. *Right:* Significant interaction between STN and genotype on head yaw caused by 40 ms stimulation (2-Way ANOVA). Left and right STN stimulation causes significant ipsiversive head yaw when compared within Cre mice, and vs. controls, for the right STN. (C) *Left:* Linear relationships between pulse width and ipsiversive head roll with left and right STN stimulation. *Right:* Significant interaction between STN and genotype on head roll caused by 40 ms stimulation (2Way ANOVA). Left and right STN stimulation causes significant ipsiversive head roll when compared within Cre mice, and vs. controls, for the left STN. (D) *Left:* Linear relationships between pulse width and upwards head pitch with left and right STN stimulation. *Right:* Significant main effect of genotype on upwards head pitch caused by 40 ms stimulation. *p<0.05, **p<0.01, ***p<0.001, ****p<0.0001. Error bars indicate SEM.

We also examined the mean latency to movement onset for each combination of kinematic variable and STN. Movements caused by 40 ms single pulses were chosen for analysis because they reliably cause movement. Body angle changes began 12.26 and 15.78 ms after left and right STN stimulation onset (two-tailed t-test: *t*_6_ = 3.561, p = 0.0119). Head yaw changes began 11.68 and 11.75 ms after left and right STN stimulation onset (two-tailed t-test: *t*_6_ = 0.03963, p = 0.9697). Head roll changes began 13.42 and 14.55 ms after left and right STN stimulation onset (two-tailed t-test: *t*_6_ = 0.8264, p = 0.4402). Head pitch changes began 12.84 and 13.16 ms after left and right STN stimulation onset (two-tailed t-test: *t*_6_ = 0.1506, p = 0.8852; Figure 5).

**Figure 5.**
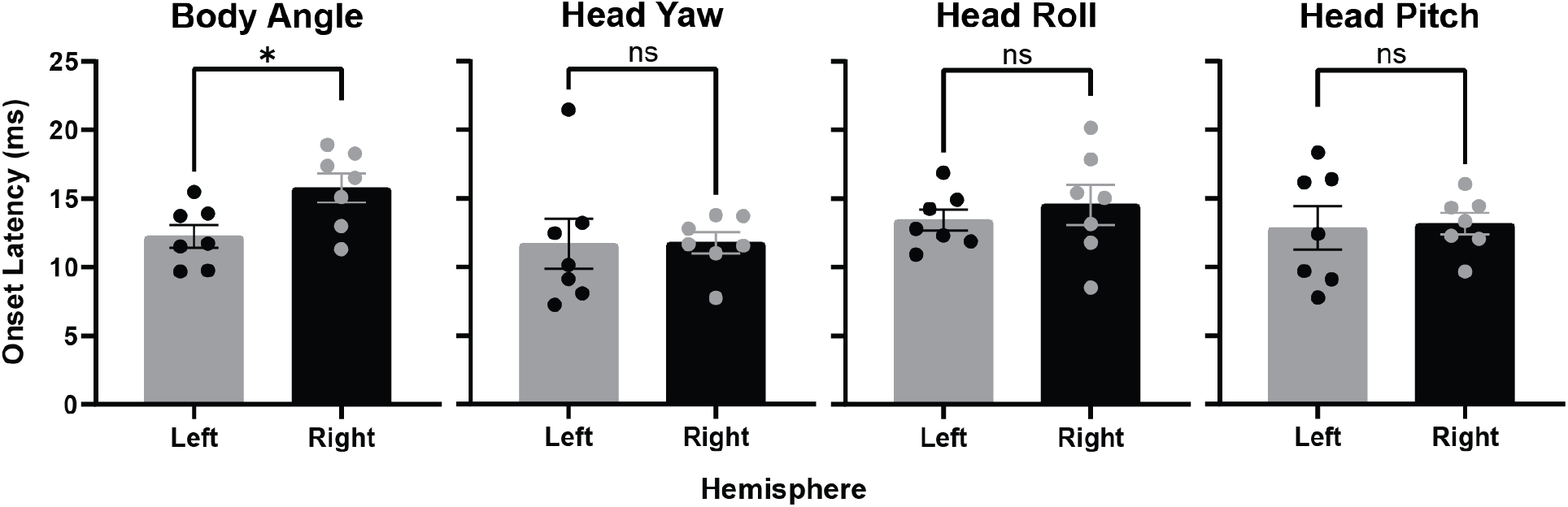
Mean latency to move from a single light pulse (40 ms). *p<0.05. Error bars indicate SEM.

We next examined mean body and head angular velocities 60 and 120 ms after single pulse stimulation onset, because velocity peaked in the ipsiversive direction after 60 ms and rebounded in the contraversive direction 60 ms later in head angle measures. At the 60 ms timepoint, linear relationships were observed between stimulation pulse width and peak effect velocity in all kinematic measures. At the 120 ms timepoint, the head velocity rebound is notable because it indicates that stimulation induced changes in yaw, roll, and pitch were quickly corrected, and thus perceived by the mice as involuntary. Consistent with the rebound being a correction of undesired movement, roll and pitch rebound velocities were linearly scaled by stimulation pulse width, and thus effect magnitude (Figure 6).

**Figure 6.**
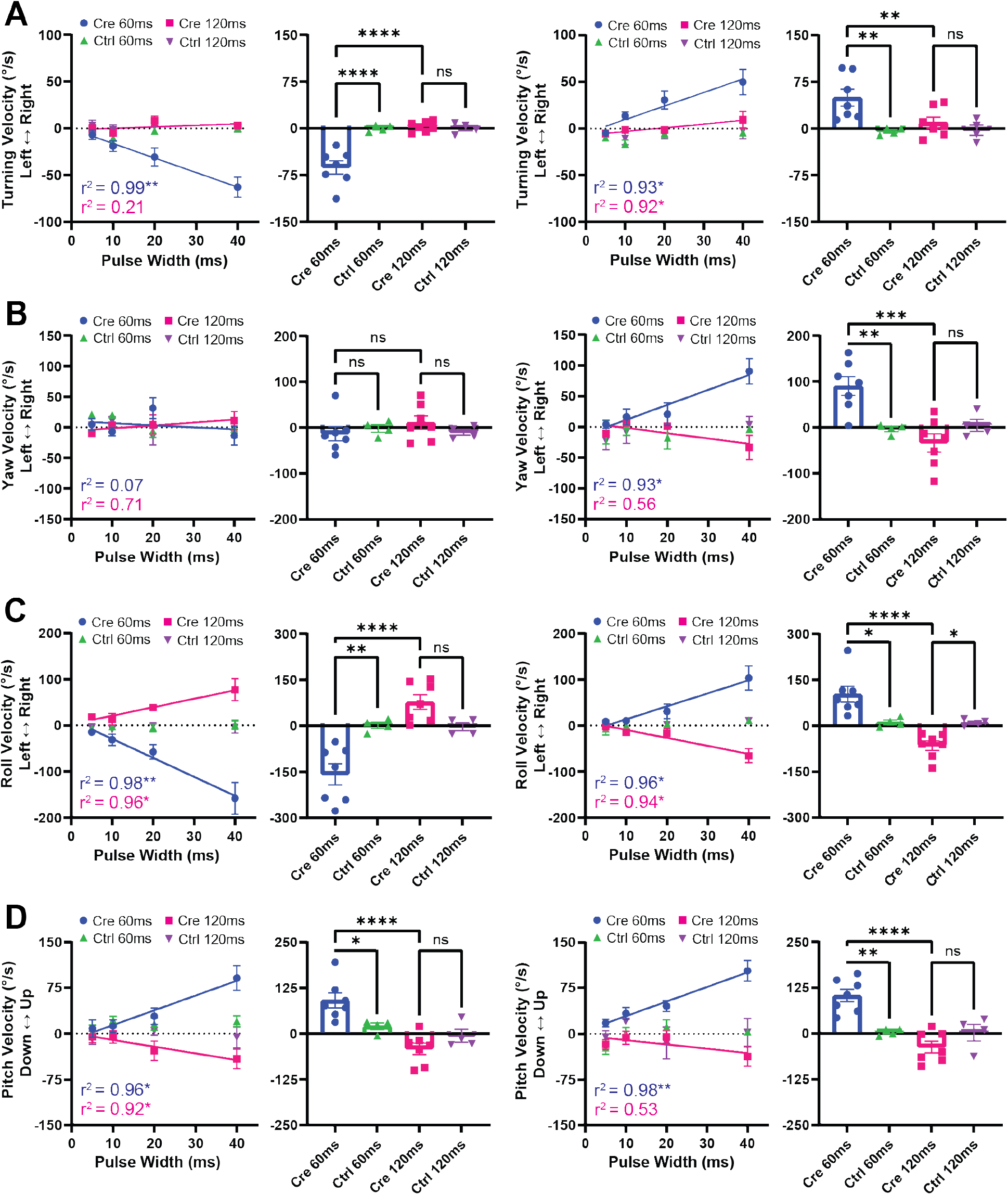
Single pulse photo-stimulation of STN VGlut2+ neurons causes velocity to peak 60ms after stimulation onset, then rebound 60 ms later in head measures. (A) Mean turning velocity. *Far Left:* Linear relationship between pulse width and ipsiversive turning velocity 60 ms after left STN stimulation. No relationship 120 ms post-stimulation. *Middle Left:* Significant interaction between timepoint and genotype on turning velocity following left STN 40 ms pulse stimulation (2-Way ANOVA). Significant ipsiversive turning velocity in Cre mice at the 60 ms timepoint. *Middle Right:* Linear relationship between pulse width and ipsiversive turning velocity 60 and 120 ms after right STN stimulation. *Far Right:* Significant interaction between timepoint and genotype on turning velocity following right STN 40 ms pulse stimulation (2-Way ANOVA). Significant ipsiversive turning velocity in Cre mice only at the 60 ms timepoint. (B) Mean head yaw velocity. *Far Left:* No relationship between pulse width and yaw velocity 60 or 120 ms after left STN stimulation. *Middle Left:* No effect of timepoint or genotype on yaw velocity following left STN 40 ms pulse stimulation (2-Way ANOVA). *Middle Right:* Linear relationship between pulse width and ipsiversive yaw velocity 60 ms after right STN stimulation. No relationship 120 ms poststimulation. *Far Right:* Significant interaction between timepoint and genotype on yaw velocity following right STN 40 ms pulse stimulation (2-Way ANOVA). Stimulation in Cre mice caused significant ipsiversive and contraversive yaw velocity at the 60 and 120 ms timepoints, respectively. (C) Mean head roll velocity. *Far Left:* Linear relationships between pulse width and ipsiversive and contraversive roll velocity 60 and 120 ms after left STN stimulation, respectively. *Middle Left:* Significant interaction between timepoint and genotype on roll velocity following left STN 40 ms pulse stimulation (2-Way ANOVA). Stimulation in Cre mice caused significant ipsiversive and contraversive roll velocity at the 60 and 120 ms timepoints, respectively. *Middle Right:* Linear relationships between pulse width and ipsiversive and contraversive roll velocity 60 and 120 ms after right STN stimulation, respectively. *Far Right:* Significant interaction between timepoint and genotype on roll velocity following right STN 40 ms pulse stimulation (2-Way ANOVA). Stimulation in Cre mice caused significant ipsiversive and contraversive roll velocity at the 60 and 120 ms timepoints, respectively. (D) Mean head pitch velocity. *Far Left:* Linear relationships between pulse width and upwards and downwards pitch velocity 60 and 120 ms after left STN stimulation, respectively. *Middle Left:* Significant interaction between timepoint and genotype on pitch velocity following left STN 40 ms pulse stimulation (2-Way ANOVA). Stimulation in Cre mice caused significant upwards and downwards pitch velocity at the 60 and 120 ms timepoints, respectively. *Middle Right:* Linear relationship between pulse width and upwards pitch velocity 60 ms after right STN stimulation. No relationship 120 ms poststimulation. *Far Right:* Significant interaction between timepoint and genotype on pitch velocity following right STN 40 ms stimulation (2-Way ANOVA). Stimulation in Cre mice caused significant upwards and downwards pitch velocity at the 60 and 120 ms timepoints, respectively. *p<0.05, **p<0.01, ***p<0.001, ****p<0.0001. Error bars indicate SEM.

In turning velocity 60 ms after unilateral stimulation onset, a linear relationship between stimulation pulse width and ipsiversive velocity was revealed in VGlut2 Cre mice (left STN linear regression: F(1,2) = 322.2, p = 0.0031, r^2^ = 0.9938; right STN linear regression: F(1,2) = 25.26, p = 0.0374, r^2^ = 0.9266). 120 ms after stimulation onset, there was no clear relationship between pulse width and turning velocity with left STN stimulation, though a statistically significant relationship was observed following right STN stimulation (left STN linear regression: p = 0.5453, r^2^ = 0.2068; right STN linear regression: p = 0.0391, r^2^ = 0.9234). There were also significant interactions between timepoint and genotype on turning velocity caused by 40 ms stimulation of left and right STN (left STN 2-Way ANOVA: F(1,9) = 19.79, p = 0.0016; right STN 2-Way ANOVA: F(1,9) = 6.169, p = 0.0348). Left STN stimulation caused significant ipsiversive turning velocity at the 60 ms timepoint versus control mice (p < 0.0001) and VGlut2-Cre mice at the 120 ms timepoint (p < 0.0001). Right STN stimulation caused significant ipsiversive turning velocity at the 60 ms timepoint versus control mice at the 60 ms timepoint (p = 0.0069) and VGlut2-Cre mice at the 120 ms timepoint (p = 0.0067; Figure 6A).

In yaw velocity 60 ms after unilateral stimulation onset, a linear relationship between stimulation pulse width and ipsiversive velocity was revealed in VGlut2 Cre mice following right, but not left, STN stimulation (left STN linear regression: F(1,2) = 0.1568, p = 0.7304, r^2^ = 0.07268; right STN linear regression: F(1,2) = 27.13, p = 0.0349, r^2^ = 0.9313). 120 ms after stimulation onset, there was no relationship between pulse width and yaw velocity with stimulation of either STN (left STN linear regression: F(1,2) = 4.927, p = 0.1566, r^2^ = 0.7113; right STN linear regression: F(1,2) = 2.519, p = 0.2534, r^2^ = 0.5575). There was, however, a significant interaction between timepoint and genotype on yaw velocity caused by right, but not left, STN 40 ms stimulation (2-Way ANOVA: F(1,18) = 10.96, p = 0.0039). Right STN stimulation caused significant ipsiversive yaw velocity at the 60 ms timepoint versus control mice at the 60 ms timepoint (p = 0.0072) and VGlut2-Cre mice at the 120 ms timepoint (p = 0.0001). Right STN stimulation also caused significant contraversive yaw velocity at the 120 versus 60 ms timepoint, indicating a rebound (p = 0.0001; Figure 6B).

In roll velocity 60 ms after unilateral stimulation onset, a linear relationship between stimulation pulse width and ipsiversive velocity was revealed in VGlut2 Cre mice (left STN linear regression: F(1,2) = 102.8, p = 0.0096, r^2^ = 0.9809; right STN linear regression: F(1,2) = 47.16, p = 0.0206, r^2^ = 0.9593). 120 ms after stimulation onset, another linear relationship between stimulation pulse width and roll velocity was observed in VGlut2 Cre mice, this time in the contraversive direction (left STN linear regression: F(1,2) = 45.44, p = 0.0213, r^2^ = 0.9578; right STN linear regression: F(1,2) = 29.90, p = 0.0319, r^2^ = 0.9373). There were also significant interactions between timepoint and genotype on roll velocity caused by left and right STN 40 ms stimulation (left STN 2-Way ANOVA: F(1,18) = 17.03, p = 0.0006; right STN 2-Way ANOVA: F(1,18) = 16.15, p = 0.0008). For the left STN, stimulation caused significant ipsiversive roll velocity at the 60 ms timepoint versus control mice at the 60 ms timepoint (p = 0.0022) and VGlut2-Cre mice at the 120 ms timepoint (p < 0.0001). Stimulation also caused significant contraversive roll velocity at the 120 versus 60 ms timepoint (p < 0.0001). For the right STN, stimulation caused significant ipsiversive roll velocity at the 60 ms timepoint versus control mice at the 60 ms timepoint (p = 0.0116) and VGlut2-Cre mice at the 120 ms timepoint (p < 0.0001). Stimulation also caused significant contraversive roll velocity at the 120 ms timepoint versus control mice at the 120 ms timepoint p = 0.0392) and VGlut2-Cre mice at the 60 ms timepoint (p < 0.0001; Figure 6C).

In measure of mean pitch velocity 60 ms after unilateral stimulation onset, a linear relationship between stimulation pulse width and upwards velocity was revealed in VGlut2 Cre mice (left STN linear regression: F(1,2) = 54.26, p = 0.0179, r^2^ = 0.9645; right STN linear regression: F(1,2) = 100.9, p = 0.0098, r^2^ = 0.9806). 120 ms after stimulation onset, left, but not right, STN stimulation revealed a linear relationship between stimulation pulse width and downwards pitch velocity (left STN linear regression: F(1,2) = 23.39, p = 0.0402, r^2^ = 0.9212; right STN linear regression: F(1,2) = 2.297, p = 0.2689, r^2^ = 0.5345). There were also significant interactions between timepoint and genotype on pitch velocity caused by left and right STN 40 ms stimulation (left STN 2-Way ANOVA: F(1,18) = 7.900, p = 0.0116; right STN 2-Way ANOVA: F(1,18) = 15.21, p = 0.0010). For the left STN, stimulation caused significant upwards pitch velocity at the 60 ms timepoint versus control mice at the 60 ms timepoint (p = 0.0320) and VGlut2-Cre mice at the 120 ms timepoint (p < 0.0001). Stimulation also caused significant downwards pitch velocity at the 120 versus 60 ms timepoint (p < 0.0001). For the right STN, stimulation caused significant upwards head pitch velocity at the 60 ms timepoint versus control mice at the 60 ms timepoint (p = 0.0019) and VGlut2-Cre mice at the 120 ms timepoint (p < 0.0001). Stimulation also caused significant downwards head pitch velocity at the 120 versus 60 ms timepoint (p < 0.0001; Figure 6D).

The head rebound is also caused by individual laser pulses in 10 Hz stimulation trains. To demonstrate this, we illustrate mean pulse driven changes in velocity and angle for all kinematic measures, across time. The rebound is clearly evident in the velocity and angular change plots for roll and pitch following stimulation of either STN, as well as for yaw following stimulation of the left STN. For yaw and roll, velocity first peaks in the direction ipsilateral to the stimulated hemisphere, then peaks again in the contralateral direction. This can be seen as an initial ipsiversive movement, followed by a contraversive correction. For pitch, velocity peaks first in the upwards direction, then peaks again in the downwards direction (Figure 7).

**Figure 7.**
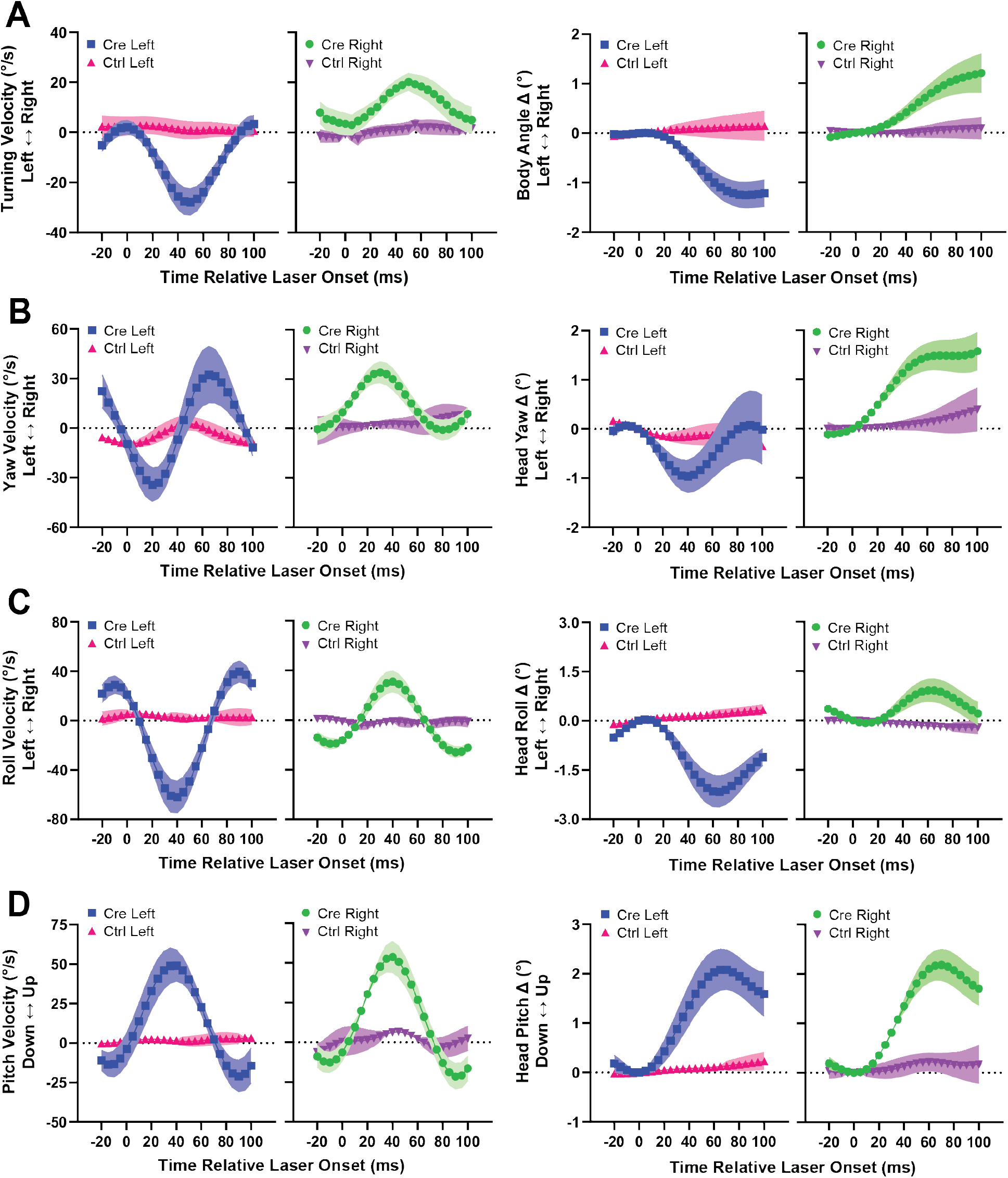
Individual laser pulses delivered to STN VGlut2+ neurons as part of 10Hz photostimulation trains cause head rebound movement. *Far Left:* Mean velocity change caused by left STN stimulation. *Middle Left:* Mean velocity change caused by right STN stimulation. *Middle Right:* Mean angular change caused by left STN stimulation. *Far Right:* Mean angular change caused by right STN stimulation. (A) Body angle. (B) Head yaw. (C) Head roll. (D) Head pitch. Error bars indicate SEM.

## Discussion

We found that selective activation of glutamatergic (vGlut2+) STN projection neurons produces systematic and highly predictable changes in head and body orientation. Unilateral stimulation consistently produces turning of the body, as well as head yaw and roll movements, in the ipsiversive direction, towards the side of stimulation. Regardless of the side of stimulation, there is also a net effect of raising the head (pitch change). These kinematic changes are a linear function of stimulation parameters (**Figures 3 & 4**). To our knowledge this is the first clear demonstration that STN output can quantitatively determine movement kinematics.

The extremely short latency from stimulation onset to movement initiation is also notable. On average movement began around 10-15 ms after stimulation onset (**Figure 5**). The short latency and the remarkably high precision of stimulation-induced movement suggest the STN output can rapidly influence effectors to alter movement trajectories. Anatomically, the STN not only projects to BG output nuclei but also midbrain and brainstem areas such as the pedunculopontine nucleus (Watson et al., 2021). It is possible that direct projections to downstream brainstem regions may be responsible for the observed effects.

Interestingly, head angle changes due to a single light pulse are often quickly corrected by the animal if the pulse is not immediately followed by another. There was often a ‘rebound’ movement in the opposite direction after 50-60 ms, bringing the head back towards the baseline position (**Figure 6**). Rebound movements are also observed when examining roll, pitch, and yaw following individual laser pulses in 10 Hz laser trains. Shortly after pulse onset, velocities peak in the expected direction, then rebound to peak again in the opposite direction before the onset of the next pulse (**Figure 7**). When stimulation frequency increases from 10 to 20 Hz, the gap between pulses shortens, so there is no time to recover between pulses. These subtle movement effects have never been reported before, as they could only be measured using our 3D motion capture system.

The quick correction of movement caused by stimulation suggests that the stimulation did not produce a volitional action, but caused an undesirable reflexive deviation that acted as a disturbance. This is supported by the short latency of the movement onset, which is more similar to reflexive movements and shorter than latencies following cortical or striatal stimulation. It is unclear how this rebound movement is generated. One possibility is that it is generated by the STN-GPe circuit, which is characterized by strong reciprocal connectivity. Normally, STN and GPe do not show correlated activity, but synchronous activation of STN output by ChR2 may have caused the GABAergic output neurons of the GPe to fire, in turn causing sharp inhibition of the STN and rebound movement (Bevan et al., 2002). This possibility remains to be tested.

Recent work in rats found GPe neurons (arkypallidal) are selectively activated on successful stop trials just before cancellation of movement-related activity in the striatum (Schmidt et al., 2013; Mallet et al., 2016; Schmidt and Berke, 2017). It was concluded that the arkypallidal neurons in the GPe project directly to the striatum to stop ongoing behavior. Higher STN activity was also associated with longer reaction times, suggesting that the STN could pause behavior. However, the only behavioral measure were beam breaks that detect nose pokes. While clearly rats never stopped breaking the beam, with such measures it is impossible to know if movements were suppressed. Alternatively, their results could be explained by STN’s role in causing ipsiversive movement and head deviation, as demonstrated in the present study.

Previous work has established that direct pathway activation produces contraversive rather than ipsiversive movements, whereas indirect pathway activation can produce ipsiversive turning movements (Kravitz et al., 2010). Via either direct excitation of SNr or projections the GPe, STN output is expected to increase output from the SNr. Our observation is therefore consistent with ipsiversive turning observed after indirect pathway activation. Many SNr neurons show high correlations with postural disturbances and head position, suggesting SNr output represents a reference signal used by downstream circuits to guide body and head position (Barter et al., 2014; Barter et al., 2015; Yin, 2017). It is well established that unilateral injections of the GABA-A antagonist picrotoxin into the SNr could produce ipsiversive turning (Olpe et al., 1977). Similarly, STN photo-stimulation could generate movement by exciting the SNr, though the very short latency of the movements evoked suggests a more direct route via direct STN projections to the brainstem.

Finally, the STN’s role in movement initiation has implications for interpreting the effects of STN DBS (Vitek, 2008). According to the standard model, dopamine depletion causes hypoactivity of direct pathway neurons and hyperactivity of indirect pathway neurons, resulting in excessive inhibition of prokinetic thalamic and brainstem targets (Albin et al., 1989; Alexander and Crutcher, 1990; Chevalier and Deniau, 1990; DeLong, 1990). It is commonly believed that STN DBS suppresses STN output, thereby making it easier for the direct pathway to disinhibit the thalamus and brainstem and restore normal movement (Chevalier and Deniau, 1990; Benazzouz et al., 1995; Benabid et al., 2001; Beurrier et al., 2001; Wichmann et al., 2018). However, it remains unclear whether STN DBS suppresses STN output, or disrupts abnormal bursting, synchronization, and oscillations in the basal ganglia (Hashimoto et al., 2003; Degos et al., 2005; Meissner et al., 2005; Shi et al., 2006; Hammond et al., 2007; Galvan and Wichmann, 2008; Hahn et al., 2008; McConnell et al., 2012; Zhuang et al., 2018; Yu et al., 2020). Recent work found that selective activation of glutamatergic projections from the parafascicular nucleus of the thalamus to the STN restores normal movement in a mouse model of PD (Watson et al., 2021). These results also support the hypothesis that that STN output can directly generate movements. Moreover, in dopamine-depleted mice with complete akinesia, the activation of the Pf-STN pathway at 25 Hz is sufficient to rescue movements. This frequency is comparable to the frequencies used in the present study, but much lower than that used in traditional STN DBS (~130 Hz). It is important to note that traditional DBS using electrical stimulation is not specific to cell bodies or a specific population of projection neurons, but likely activate a combination of afferent terminals from multiple areas as well as fibers of passage. The clinical efficacy of DBS could be due to the activation, rather than suppression, of STN neurons.

In short, using precise targeting of STN output neurons, we were able to manipulate STN output and at the same time quantify evoked movements with high temporal and spatial resolution. We covered a highly linear relationship between stimulation parameters and kinematics, suggesting that STN can quantitatively scale movements, especially turning in the ipsiversive direction. These results question the prevailing model that STN functions to suppress movements, and suggests instead that STN output plays an active role in scaling movement parameters.

## Acknowledgments

We would like to thank Konstantin Bakhurin for help with data analysis, Guozhong Yu for help with surgery and histology, and Francesco Ulloa Severino for help with Adobe Illustrator. This work was supported by NIH grants NS094754 and MH112883 to H.H.Y.

## References

Albin RL, Young AB, Penney JB (1989) The functional anatomy of basal ganglia disorders. Trends Neurosci 12:366–375.

Alexander GE, Crutcher MD (1990) Functional architecture of basal ganglia circuits: neural substrates of parallel processing. Trends Neurosci 13:266–271.

Aron AR, Poldrack RA (2006) Cortical and subcortical contributions to Stop signal response inhibition: role of the subthalamic nucleus. J Neurosci 26:2424–2433.

Barter JW, Castro S, Sukharnikova T, Rossi MA, Yin HH (2014) The role of the substantia nigra in posture control. Eur J Neurosci 39:1465–1473.

Barter JW, Li S, Sukharnikova T, Rossi MA, Bartholomew RA, Yin HH (2015) Basal ganglia outputs map instantaneous position coordinates during behavior. J Neurosci 35:2703–2716.

Bartholomew RA, Li H, Gaidis EJ, Stackmann M, Shoemaker CT, Rossi MA, Yin HH (2016) Striatonigral control of movement velocity in mice. The European journal of neuroscience 43:1097–1110.

Baunez C, Robbins TW (1997) Bilateral lesions of the subthalamic nucleus induce multiple deficits in an attentional task in rats. Eur J Neurosci 9:2086–2099.

Baunez C, Nieoullon A, Amalric M (1995) In a rat model of parkinsonism, lesions of the subthalamic nucleus reverse increases of reaction time but induce a dramatic premature responding deficit. J Neurosci 15:6531–6541.

Benabid AL, Koudsie A, Benazzouz A, Vercueil L, Fraix V, Chabardes S, Lebas JF, Pollak P (2001) Deep brain stimulation of the corpus luysi (subthalamic nucleus) and other targets in Parkinson’s disease. Extension to new indications such as dystonia and epilepsy. J Neurol 248 Suppl 3:III37–47.

Benazzouz A, Piallat B, Pollak P, Benabid AL (1995) Responses of substantia nigra pars reticulata and globus pallidus complex to high frequency stimulation of the subthalamic nucleus in rats: electrophysiological data. Neurosci Lett 189:77–80.

Beurrier C, Bioulac B, Audin J, Hammond C (2001) High-frequency stimulation produces a transient blockade of voltage-gated currents in subthalamic neurons. J Neurophysiol 85:1351–1356.

Bevan MD, Magill PJ, Terman D, Bolam JP, Wilson CJ (2002) Move to the rhythm: oscillations in the subthalamic nucleus-external globus pallidus network. Trends Neurosci 25:525–531.

Chevalier G, Deniau JM (1990) Disinhibition as a basic process in the expression of striatal functions. Trends Neurosci 13:277–280.

Degos B, Deniau JM, Thierry AM, Glowinski J, Pezard L, Maurice N (2005) Neuroleptic-induced catalepsy: electrophysiological mechanisms of functional recovery induced by high-frequency stimulation of the subthalamic nucleus. J Neurosci 25:7687–7696.

DeLong MR (1990) Primate models of movement disorders of basal ganglia origin. Trends Neurosci 13:281–285.

DeLong MR, Wichmann T (2001) Deep brain stimulation for Parkinson’s disease. Ann Neurol 49:142–143.

Eagle DM, Baunez C, Hutcheson DM, Lehmann O, Shah AP, Robbins TW (2008) Stop-signal reaction-time task performance: role of prefrontal cortex and subthalamic nucleus. Cereb Cortex 18:178–188.

Fife KH, Gutierrez-Reed NA, Zell V, Bailly J, Lewis CM, Aron AR, Hnasko TS (2017) Causal role for the subthalamic nucleus in interrupting behavior. Elife 6.

Galvan A, Wichmann T (2008) Pathophysiology of parkinsonism. Clin Neurophysiol 119:1459–1474.

Guillaumin A, Serra GP, Georges F, Wallen-Mackenzie A (2021) Experimental investigation into the role of the subthalamic nucleus (STN) in motor control using optogenetics in mice. Brain Res 1755:147226.

Hahn PJ, Russo GS, Hashimoto T, Miocinovic S, Xu W, McIntyre CC, Vitek JL (2008) Pallidal burst activity during therapeutic deep brain stimulation. Exp Neurol 211:243–251.

Hammond C, Bergman H, Brown P (2007) Pathological synchronization in Parkinson’s disease: networks, models and treatments. Trends Neurosci 30:357–364.

Hashimoto T, Elder CM, Okun MS, Patrick SK, Vitek JL (2003) Stimulation of the subthalamic nucleus changes the firing pattern of pallidal neurons. J Neurosci 23:1916–1923.

Heston J, Friedman A, Baqai M, Bavafa N, Aron AR, Hnasko TS (2020) Activation of Subthalamic Nucleus Stop Circuit Disrupts Cognitive Performance. eNeuro 7.

Kravitz AV, Freeze BS, Parker PR, Kay K, Thwin MT, Deisseroth K, Kreitzer AC (2010) Regulation of parkinsonian motor behaviours by optogenetic control of basal ganglia circuitry. Nature 466:622–626.

Mallet N, Schmidt R, Leventhal D, Chen F, Amer N, Boraud T, Berke JD (2016) Arkypallidal Cells Send a Stop Signal to Striatum. Neuron 89:308–316.

McConnell GC, So RQ, Hilliard JD, Lopomo P, Grill WM (2012) Effective deep brain stimulation suppresses low-frequency network oscillations in the basal ganglia by regularizing neural firing patterns. J Neurosci 32:15657–15668.

Meissner W, Leblois A, Hansel D, Bioulac B, Gross CE, Benazzouz A, Boraud T (2005) Subthalamic high frequency stimulation resets subthalamic firing and reduces abnormal oscillations. Brain 128:2372–2382.

Nambu A, Tokuno H, Takada M (2002) Functional significance of the cortico-subthalamo-pallidal ‘hyperdirect’ pathway. Neurosci Res 43:111–117.

Olpe H-R, Schellenberg H, Koella W (1977) Rotational behavior induced in rats by intranigral application of GABA-related drugs and GABA antagonists. European journal of pharmacology 45:291–294.

Parent A, Hazrati LN (1995) Functional anatomy of the basal ganglia. II. The place of subthalamic nucleus and external pallidum in basal ganglia circuitry. Brain research Brain research reviews 20:128–154.

Schmidt R, Berke JD (2017) A Pause-then-Cancel model of stopping: evidence from basal ganglia neurophysiology. Philos Trans R Soc Lond B Biol Sci 372.

Schmidt R, Leventhal DK, Mallet N, Chen F, Berke JD (2013) Canceling actions involves a race between basal ganglia pathways. Nat Neurosci 16:1118–1124.

Shi LH, Luo F, Woodward DJ, Chang JY (2006) Basal ganglia neural responses during behaviorally effective deep brain stimulation of the subthalamic nucleus in rats performing a treadmill locomotion test. Synapse 59:445–457.

Smith Y, Bolam JP (1989) Neurons of the substantia nigra reticulata receive a dense GABA-containing input from the globus pallidus in the rat. Brain Res 493:160–167.

Uslaner JM, Robinson TE (2006) Subthalamic nucleus lesions increase impulsive action and decrease impulsive choice - mediation by enhanced incentive motivation? Eur J Neurosci 24:2345–2354.

Vitek JL (2008) Deep brain stimulation: how does it work? Cleve Clin J Med 75 Suppl 2:S59–65.

Watson GD, Hughes RN, Petter EA, Fallon IP, Kim N, Severino FPU, Yin HH (2021) Thalamic projections to the subthalamic nucleus contribute to movement initiation and rescue of parkinsonian symptoms. Science Advances 7:eabe9192.

Wichmann T, Bergman H, DeLong MR (2018) Basal ganglia, movement disorders and deep brain stimulation: advances made through non-human primate research. J Neural Transm (Vienna) 125:419–430.

Yin HH (2017) The Basal Ganglia in Action. Neuroscientist 23:299–313.

Yu C, Cassar IR, Sambangi J, Grill WM (2020) Frequency-Specific Optogenetic Deep Brain Stimulation of Subthalamic Nucleus Improves Parkinsonian Motor Behaviors. J Neurosci 40:4323–4334.

Zhuang QX, Li GY, Li B, Zhang CZ, Zhang XY, Xi K, Li HZ, Wang JJ, Zhu JN (2018) Regularizing firing patterns of rat subthalamic neurons ameliorates parkinsonian motor deficits. J Clin Invest 128:5413–5427.

